# Brain organization, not size alone, as key to high-level vision: Evidence from marmoset monkeys

**DOI:** 10.1101/2020.10.19.345561

**Authors:** Alexander J.E. Kell, Sophie L. Bokor, You-Nah Jeon, Tahereh Toosi, Elias B. Issa

## Abstract

Bigger brains are thought to support richer abilities, including perceptual abilities. But bigger brains are typically organized differently (e.g., with more cortical areas). Thus, the extent to which a neural system’s size versus organization underlies complex abilities remains unclear. The marmoset monkey is evolutionarily peculiar: it has a small brain, yet many cortical areas. We used this natural experiment to test organization as source of high-level visual abilities independent of size, via large-scale psychophysics comparing marmosets to different species on identical tasks. Marmosets far out—performed rats—a marmoset-sized rodent—on a simple visual recognition task. On another visual task, which is difficult for both humans and machines, marmosets achieved high performance. Strikingly, their image-by-image behavior revealed that they did so in a manner highly similar to humans—marmosets were nearly as human-like as were macaques. These results suggest a key role for brain organization—not simply size—in the evolution of sophisticated abilities.

## Introduction

A bigger brain is thought to confer an organism with richer abilities^1–6^. Originally, a large brain relative to body size—a high “encephalization quotient"^1^—was believed to be critical, but recent work has suggested absolute size may be decisive^3–5^, and perhaps the number of cortical neurons in particular^6–8^. While brain size is typically studied in relation to domains considered “cognitive“^2–6^, the hypothesis is thought to apply generically, including to high-level perceptual abilities^9–11^.

Bigger brains, however, tend to be organized differently than smaller brains. As brain size increases, each neuron synapses with a smaller proportion of total neurons, resulting in a more modular organization (e.g., with more cortical areas)^11–13^. Organizational factors like these have also been hypothesized to bestow richer abilities^11,13,14^, but the correlation between brain size and organization has made it difficult to assess the unique contribution of brain organization.

The marmoset monkey has a small brain yet many cortical areas, and therefore offers a natural experiment to address this question. Weighing just 350 grams, marmosets are phyletic dwarves^15–17^, shrunken from far larger ancestors. Phyletic dwarfism is evolutionarily unusual^18,19^ and can lead to atypical configurations of phenotypes^15,20^. Whether a legacy of this evolutionary history or not, the marmoset brain has many cortical areas (~120)^21–23^—by comparison, mice, macaques, and humans respectively have ~40, ~140 and ~180^23^. Most animals, including larger primates such as squirrel monkeys, owl monkeys, and rhesus macaques, follow a lawful relationship between the size of the cortical sheet and the number of cortical areas, but marmosets have far more areas than expected given their small brain^24^.

We took advantage of the unusual biology of the marmoset monkey to test the role of brain size versus organization in supporting high-level visual abilities. Despite its apparent ease to humans, invariantly recognizing objects is a highly sophisticated behavior, the culmination of many stages of complex visual processing^25–27^. Indeed, human levels of performance on such high-level visual tasks is not supported by early or even intermediate stages of visual cortex^26,28^, and engineering systems have only recently been able to achieve human levels of performance^29,30^.

The marmoset brain has organizational features believed to be critical for these visual behaviors, including a large number of hierarchically organized ventral visual areas^31,32^. But a marmoset’s brain is two orders of magnitude smaller than a human’s (8 vs 1500 grams), with orders of magnitude fewer cortical neurons, both in aggregate and per cortical area. These factors are thought to lead to lesser abilities in general^10,11,14^, and have been suggested to limit the marmosets’ high-level abilities in particular^33,34^. Here we tested whether marmosets can achieve high performance on sophisticated visual tasks, first comparing their behavior with that of rats—a marmoset-sized rodent but with fewer cortical areas—as a test of the effect of varying brain organization and then with macaques and humans, larger-brained but with roughly similar numbers of cortical areas. lf brain size is of primary importance, then we would expect marmoset visual behavior to be much closer to the similarly sized rat and far from the human. Alternatively, if brain organization plays an important role, we expect marmosets’ behavior to be closer to that of humans. Critically, it’s unclear how adequate, if at all, this similarity in organization is for closing the gap between a small animal and the human.

## Results

### Marmosets far outperformed rats on a simple visual recognition task

We began by comparing marmoset visual behavior to that of rats, a marmoset-sized rodent that has a modestly smaller brain, but far fewer cortical areas^35^. Rats and other rodents (e.g., mice) have become increasingly popular choice for studying vision in the field of neuroscience^36–38^.

We tested marmosets on a visual task that had been first used in rats, the results of which had been interpreted as evidence of the rodent’s sophisticated visual abilities^36^. Animals were trained on a two-alternative forced-choice task to recognize two synthetic objects in isolation on black backgrounds and then tested on novel images of those same isolated objects under previously unencountered conjunctions of rotations and scales (Fig. 1a). Using identical images and task design, we found that marmosets performed substantially better than rats (mean accuracy marmosets: 93%, rats: 71%; two-tailed t test: t_53_ = 18.4, p = 1.14 × 10^−24^; Fig. 1b and 1c). We then examined performance at a finer scale, separating out those images that marmosets and rats had encountered in training and those that they had not. When encountering novel rotations and scales, rats performed far worse, whereas the marmosets were unaffected and instead simply invariant (Fig. 1d and 1e; ANOVA: species-by-scale interaction F_5,78_ = 39.9, p = 3.81 × 10^−20^, species-by-rotation interaction F_8,78_ = 7.62, p = 1.78 × 10^−7^). These differences were observed even though image size was scaled up to 40 degrees for the rats, in order to accommodate their coarse visual acuity (visual acuity of human, marmoset, and rat, respectively: 60, 30, and 1 cycles per degree^39–41^). These results show that marmosets’ visual abilities substantially surpass those of an animal with a relatively comparable brain size but fewer cortical areas.

**Fig. 1.**
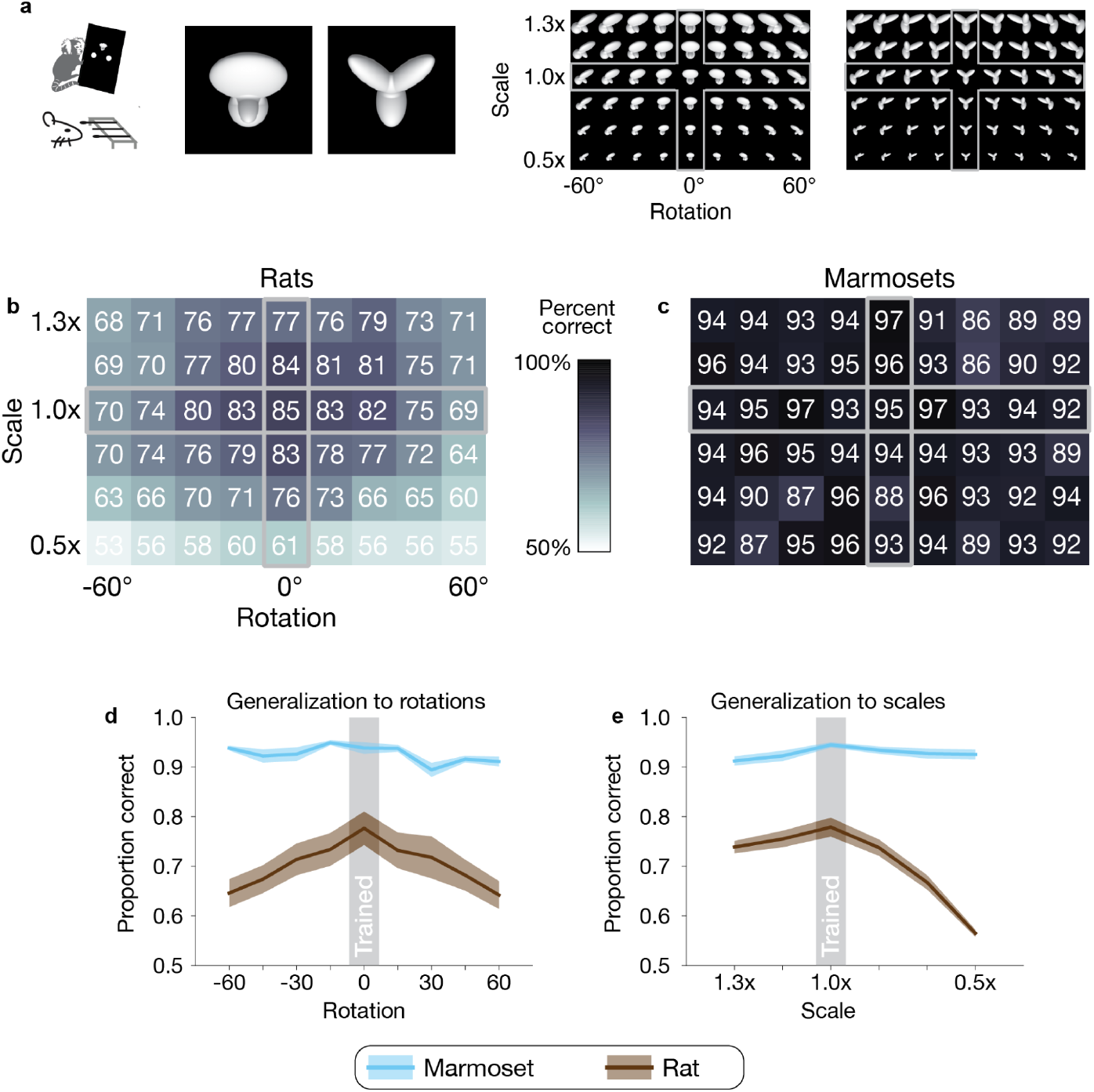
Comparing marmosets to rats, a marmoset-sized rodent. **(a)** The two objects used in a prior rodent study^36^ (left) and all images generated by varying object rotation and scale (right). To test generalization, rats and marmosets were trained on a subset of images (the cross outlined in gray) and then evaluated on all. **(b)** Rat accuracy at each rotation and scale, reproduced from Zoccolan et al., 2009. Overlaid: Percent correct for each image. Gray outline indicates images on which marmosets and rats were trained. **(c)** Marmoset accuracy at each rotation and scale. Plotting conventions, including color scale, same as (b). **(d)** Generalization to novel images at each rotation. Both species were trained at 0° (frontal view). Error bars are SEM over scales. **(e)** Generalization to novel images at each scale. Both species were trained at 1.0x. Error bars are SEM over rotations.

In a subsequent control experiment, however, we found that this task, at least from one perspective, is not particularly high level. A linear classifier trained simply on the image pixels achieved an accuracy of 97.5%, demonstrating that high task performance could be attained with retina-like representations (i.e., image pixels). By contrast, the strongest form of invariant object recognition is thought to require multiple stages of complex cortical processing^26,27,42^—indeed, in the brain, high performance cannot be achieved until after multiple stages of cortical processing^26^. As this first task appears not to be a strong test of the marmosets’ high-level abilities, we tested marmosets on a far more challenging visual task below.

### Marmosets achieved high performance on a challenging high-level visual task

We then evaluated marmosets’ visual behavior on a task that used naturalistic images and intuitively appears far more challenging than the task used in rodents above: the invariant identification of objects in the face of changes in object scale, position, and pose on rich, natural backgrounds (Fig. 2a). On each trial, participants (human or marmoset) chose which of two basic-level objects (e.g., camel or wrench) was present in a briefly flashed image (Fig. 2b, Supp. Fig. 1).

**Figure 2.**
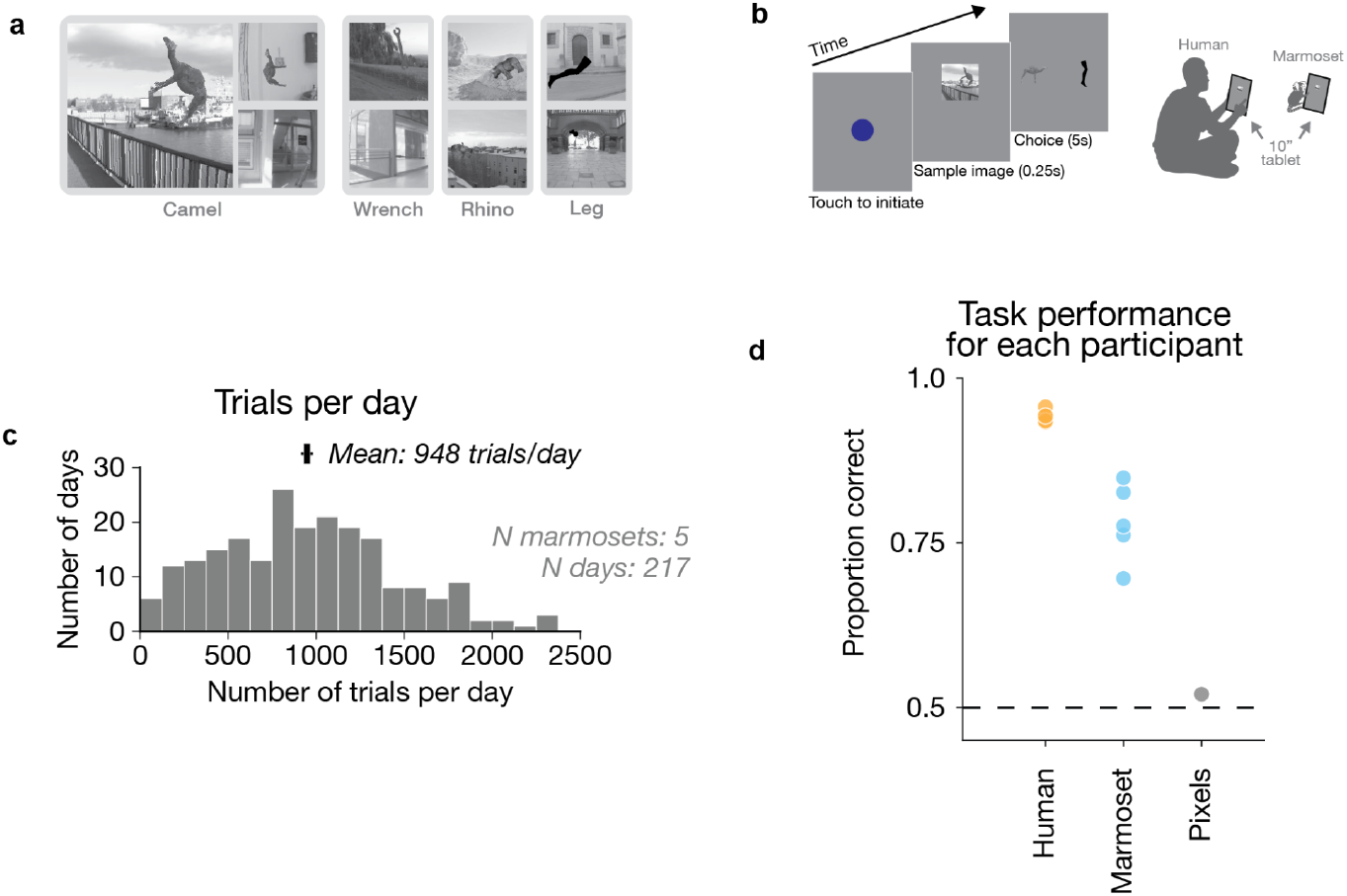
High-level invariant object recognition task and marmoset behavior. **(a)** Example images (of 400 total) from the four objects tested. **(b)** Task design. **(c)** Histogram of trials per day by marmosets. **(d)** Task performance for humans, each marmoset subject, and a baseline control model. While performance was measured on identical images, presentation time differed across marmosets and humans (250 and 50 msec, respectively).

We first tested whether this task was indeed challenging, evaluating human performance on these images (n=7; n trials=146,496). Consistent with previous work that used identical images^43^, we found that humans performed this two-alternative forced-choice task well, but by no means perfectly, achieving 94% accuracy (Fig. 2d). Furthermore, we found that a linear classifier trained on pixels for this task was near chance performance (50%), suggesting that this task required nontrivial visual processing. Taken together, these results show that these images comprise a challenging object recognition task and are therefore a strong test of a high-level complex visual ability.

We then tested marmosets’ abilities on this challenging high-level perceptual task. We evaluated marmosets on the same 400 images on the same touchscreen tablets (n=5; n trials=205,667; Fig. 2c; Supp. Fig. 2, Supp. Video 1). Remarkably, marmoset monkeys performed at 80% accuracy. Mean accuracy for each marmoset was 88%, 86%, 78%, 76%, and 70% (Fig. 2d). We employed no subject inclusion or selection criteria—and thus these individual performances are representative of marmosets’ capabilities, rather than a reflection of a few outlier high-performing animals. The high performance of each subject demonstrates that this small-brained New World primate performs well at a demanding high-level perceptual task.

### Marmosets exhibited human-like error patterns

While marmosets and humans both achieved high performance on this task, it remained unclear the extent to which the two used similar strategies. Marmosets are small, arboreal primates with substantially different visual ecology and affordances than humans, and thus it is not obvious that their visual strategies would be similar to those of humans. Moreover, while the human participants in this study could be instructed as to the task design, the marmoset monkeys had to be trained with operant conditioning to perform this task over weeks (note: they were trained with different images than used for testing). Perhaps through this use of rewards we trained in a non-ecological behavior and, as a consequence, marmosets—while high achieving—are performing this task in a starkly different manner than the untrained humans. By contrast, common strategies across humans and marmosets, in spite of training differences, could be one consequence of common aspects of brain organization across the two species.

We tested the performance signatures of marmosets and humans in this task by comparing image-by-image performance across species, under the logic that a similar visual strategy would result in similar images being easy or difficult for both marmosets and humans. We computed the difficulty of each image and subtracted off the average difficulty of each object (Fig. 3a; see Methods for details). Unlike one-dimensional summaries of overall performance, this image-by-image 400-length vector is a rich, high-dimensional signature of visual perception that is robust to global, non-perceptual factors like attentional lapses or motor errors. Such global factors lead to absolute shifts in performance but leave the relative image-by-image pattern intact. This metric was therefore well suited for comparing marmosets and humans despite their differences in mean performance.

**Fig. 3.**
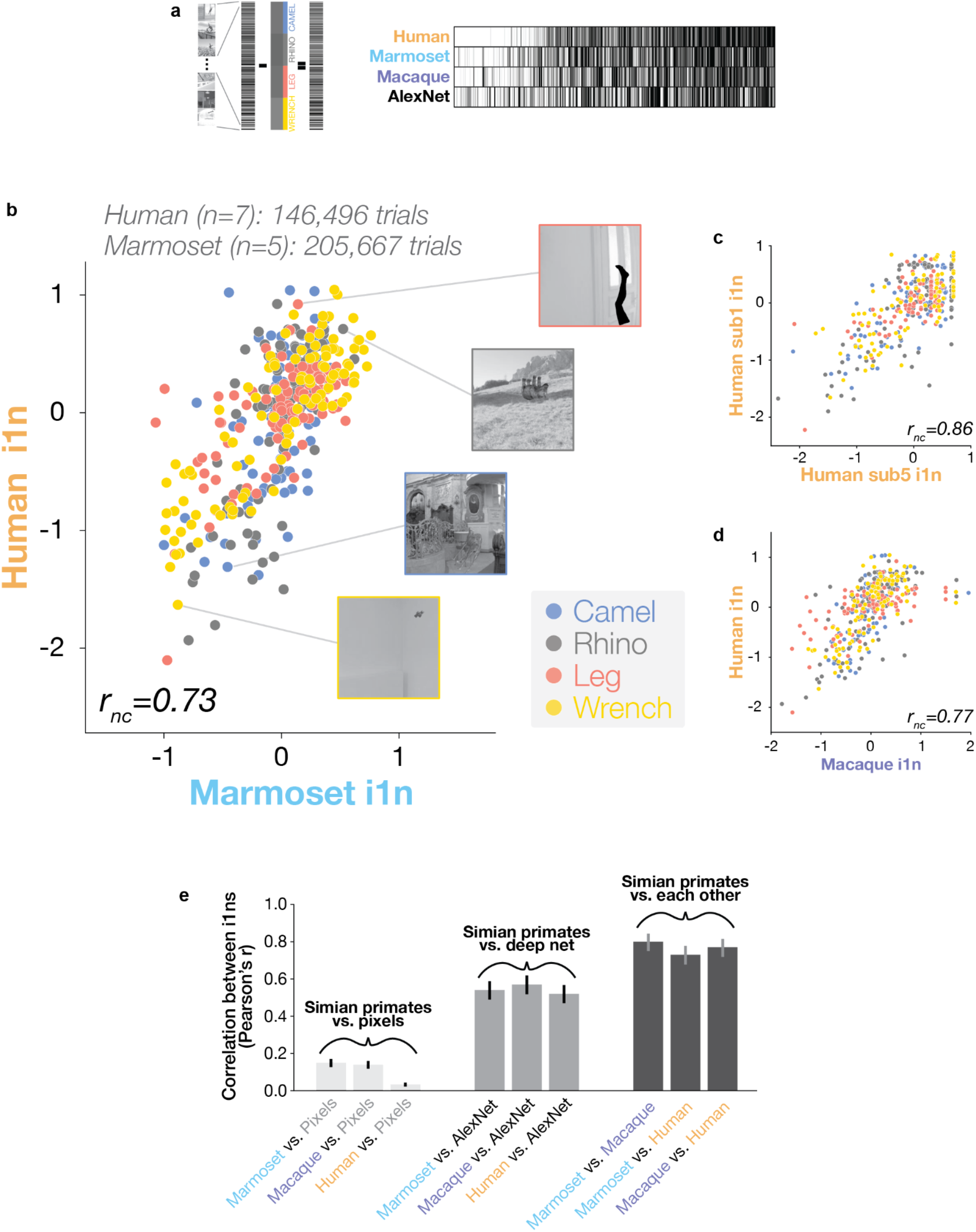
Image-by-image behavioral comparisons of simian primates. **(a)** Image-by-image behavioral metric (i1n). Left: Normalization subtracts off object-level means, leaving the fine-grained, image-by-image pattern. Right: i1n signatures (barcodes) for marmosets, humans, macaques, and an artificial deep neural network (AlexNet), sorted by left-out human data. **(b)** Scatter comparing marmoset and human behavior for each of 400 images. Color indicates object; *r_nc_* denotes noise-corrected correlation (correcting for test-retest reliability of the data). Inset: Images for example points. **(c)** Scatter of two human participants’ against each other. **(d)** Scatter comparing macaque and human behavior. **(e)** Pairwise correlation between simian primates’ i1n performance signatures and those of pixel classifiers, a deep neural network, and each other. Error bars indicate 95% confidence intervals.

We found that marmosets and humans tended to find the same images easy or difficult, as image-by-image performance was remarkably correlated (r = 0.73, p = 8.91 × 10^−68^; Fig. 3b). Humans have approximately twice the foveal acuity as marmosets (~60 vs. 30 cycles per degree^39,40^), and so it is not obvious how to equate image size across the species. We therefore tested marmosets at image sizes that differed by an octave, and found that marmosets exhibited human-like behavior irrespective of size (marmosets ~22° images, n trials = 98,378, r = 0.65, p = 2.17 × 10^−49^; marmosets ~11° images, n trials = 107,289: r = 0.71, p = 1.35 × 10^−62^). The high correlation between marmosets and humans is rendered even more impressive by the fact that different humans exhibited slightly different image performance signatures, even after accounting for measurement noise (mean cross-subject correlation, r = 0.88; Fig. 3c). These results suggest that marmosets—despite having both a brain and a body size 11200^th^ the mass of humans’—perform invariant visual object recognition in a strikingly human-like manner.

### Marmosets are nearly as human-like as are macaques

To better isolate the effect of brain size, we can compare marmosets to rhesus macaques since those two species have similar number of brain areas (~120 and ~140, respectively^23^) but marmosets’ brains are far smaller. Using previously collected macaque data on the same images^43^, we found that marmosets and macaques exhibited highly similar image-by-image behavior (r = 0.80, p = 2.52 × 10^−90^; Fig. 3d) and that, remarkably, macaques were only slightly more human-like than were marmosets (macaque-human r = 0.77; marmoset-human r = 0.73, dependent test of correlation coefficients^44^: t_397_ = 2.06, p = 0.040).

However, some details of experimental design in the previous work differed from ours (e.g., whether objects were interleaved within or across sessions; see Methods for details), which raises the possibility that we may have underestimated the macaque-human similarity relative to the marmoset-human similarity, despite the use of identical images across all three species. lt seems unlikely that these differences were particularly consequential for the image performance signatures. Human signatures collected across the two settings were highly correlated (r = 0.90, p = 1.32 × 10^−145^). Moreover, the macaque-human similarity reported in the previous work, when macaques and humans were in more similar task settings, was highly similar to what we estimated here across task settings (previous work^43^, macaque-human r = 0.77; here, r = 0.77). The image performance signatures therefore appeared to be primarily determined by the perceptual challenge of core basic-level object recognition, which was common across experimental settings in macaques, marmosets, and humans. Taken together, these results demonstrate how strikingly similar marmoset high-level visual behavior is to that of humans.

Does this similar image-by-image performance across visual systems of two primate species reveal common processing strategies for those two systems? Or might they instead reflect something about the images—e.g., might some images simply be intrinsically more difficult than others? lf the latter were the case, then any high-performing visual system would exhibit similar image-by-image performance^45^. To test this possibility, we evaluated the performance characteristics of high-performing artificial neural networks. We found that these networks performed this task as well as marmosets but nonetheless exhibited image-by-image performance that was substantially less human-like than marmosets’ performance (Fig. 3e; Supp. Fig. 3 and 4; AlexNet^29^-humans r = 0.52, dependent test of correlation coefficients t_397_ = 6.37, p = 5.29 × 10^−10^; see Methods for details). This result dovetails with prior work showing a gap between models and humans^43^, suggesting that the similarity between marmoset and human behavior is not a trivial consequence of high task performance, but instead may reflect a consequence of common organizational principles across the brains of the two species.

## Discussion

To dissociate the relative contribution of brain size and brain organization to high-level visual abilities, we took advantage of the natural experiment of the marmoset monkey, which has a small brain yet many cortical areas. We conducted large-scale, cross-species psychophysics comparing the visual behavior of marmosets with that of rats, macaques, and humans on identical stimuli and tasks. We found that marmosets dramatically out-performed the rodents-while rats were substantially affected by object size and pose, marmosets were simply invariant. We then compared marmosets to a primate with a brain two-hundreds times the size: humans. On an invariant object recognition task that challenges humans and machine vision systems, marmosets achieved high performance. Remarkably, marmosets’ fine-grained, image-by-image performance signature across hundreds of images closely mirrored those of humans-indeed, marmosets were nearly as human-like as were macaques. These results show that the small marmoset monkey exhibits computationally challenging high-level perceptual abilities, showing that brain organization, not size alone, plays a key role in supporting sophisticated abilities.

### High-level vision as a test domain for size versus organization

One potential concern with using a visual task is that simian primates, such as marmosets, humans, and macaques, have peripheral adaptations that increase the fidelity of the visual sensors—e.g., a high-acuity fovea with minimal retinal summation^39^. However, the neurophysiology supporting high-level visual behaviors is well established: many stages of cortical processing are required to achieve high performance on this task^25–27^, and such performance cannot be supported by even complex mid-level cortical areas^26,28^. Thus, increased sensory precision in the periphery is, by itself, insufficient to account for marmosets’ high performance, which depends on central cortical processing.

Indeed, high-level vision was an appealing test domain precisely because the underlying neuroscience is relatively advanced. Previous work made clear that marmoset brains exhibit anatomical features thought to be important for these kinds of high-level visual behaviors, including an elaboration of many visual areas, organized hierarchically^31,32^. Despite these organizational properties, it was not obvious that marmosets would close the brain size gap and be able to exhibit human-like high-level visual behaviors, as previous work had only shown similarities for relatively lower-level behaviors, such as eye movements^46,47^. Moreover, marmosets’ high-level abilities have been doubted specifically because of their small brain^33,34^. Our results reveal that marmosets are indeed capable of sophisticated, high-level visual behavior, suggesting that aspects of brain organization are important for supporting sophisticated behaviors.

### Comparing fewer species with greater precision

Previous tests of the relationship between brain size and behavioral capabilities often summarized the behavior of an animal with a single number (e.g., mean performance on a given task)^3–6^. We take a different approach: narrower in that we only examine a few species, but deeper in that we quantify the behavioral phenotype in great detail—i.e., using a 400-dimensional behavioral signature to compare marmosets, humans, and macaques. Such higher dimensional measures avoid reducing the complexities of an animal’s ethology to a single number. In so doing, they better capture the richness of the behavioral phenotype, allowing quantitative rigor without sacrificing holism or nuance. The cost of such higher-dimensional comparisons is that reliably measuring these signatures required hundreds of thousands of trials, which made it practical only to compare a handful of species—and encouraged the selection of species that offer a natural experiment, like the marmoset.

Future work could further clarify the role of brain size and organization in high-level visual abilities by using the same stimuli and tasks to assess diurnal species that likely have fewer cortical areas than the marmoset but are similarly small in size, such as lemurs or tree shrews, as well as testing non-primates like cats. Future work could also marry such higher-dimensional behavior phenotyping with higher-dimensional neural phenotyping to establish more nuanced relationships between brain structure and behavior.

### A neural system’s size, organization, and capabilities

Both brain size and organization likely matter for an animal’s abilities. Brains are strikingly metabolically expensive and their size appears to be constrained by this expense^48,49^. Consequently, organizing neural systems effectively may be a natural means to make the most of a given number of neurons. Moreover, the reported correlations between brain size and behavioral abilities are somewhat modest^3–6^, and thus leave room for additional explanatory factors, like organization, to contribute.

Brain organization has also been invoked to account for the abilities of other animals, such as birds. Despite their small brains, birds can exhibit sophisticated behavior^50,51^, which has been suggested to result from the sheer number of neurons densely packed into their forebrain^8^. Recent work, however, has also suggested that the pallium, a part of their forebrain, is organized in a rather sophisticated way, exhibiting a cortex-like microarchitecture^52^ and following information processing principles similar to those observed in cortex^53^. These similarities are at a different scale than what we focus on here (microarchitecture rather than mesoarchitecture), but nonetheless are consistent with a role of organization, not size alone, in generating an organism’s sophisticated abilities.

What aspects of brain organization support more sophisticated abilities? The organization of the marmosets’ ventral visual stream suggests, among others, two distinct factors: a greater degree of modularity (i.e., greater number of areas) and the organization of those areas into a hierarchy. Arealization may be beneficial as it may allow for greater division of labor, with different areas specializing in different functions^14^. Meanwhile, a deeper hierarchy may provide greater efficiency-indeed, theoretical work around the renaissance in artificial neural networks has shown that deeper hierarchies may reduce the number of neurons required to support a given task and the amount of data needed to learn^30,54^. Even in invertebrates such as insects, these two factors, modularity and hierarchy, have been hypothesized to account for comparatively sophisticated motor, social, and cognition abilities^55^. Future comparative work across species and anatomical characterization of their mesoarchitecture and microarchitecture can further clarify the relationship between brain organization and behavioral ability and, in doing so, help elucidate what properties of organization support rich abilities.

## Methods

### Subjects

Five common marmosets (*Callithrix jacchus*) and seven humans participated in our experiments. The five authors were among the human participants. The human data were collected in accordance with the Institutional Review Board of Columbia University Medical Center, and the marmoset data were collected in accordance with the NIH guidelines and approved by the Columbia University Institutional Animal Care and Use Committee (IACUC). We also used data from two previously published studies^36,43^ from five macaques, 1,481 humans, and six rats. These data were collected in accordance with NIH guidelines, the Massachusetts Institute of Technology Committee on Animal Care, The Massachusetts Institute of Technology Committee on the Use of Humans as Experimental Subjects (COUHES), and the Harvard Institutional Animal Care and Use Committee.

### Two-way stimulus-response task

Marmoset, human, and macaque performance was measured on the identical set of images (same object, pose, position, scale, and background). See Supplemental Figure 1 for these 400 images. Humans, macaques, and marmosets initiated each trial by touching a dot at the center of the screen. A sample image was then flashed (for marmosets 250 msec and for humans 50 msec; we varied presentation duration to induce a sufficient number of errors to yield reliable human image-by-image performance scores; the macaque data that was collected in previous work presented images for 100 msec). After the image disappeared, two example token images were presented (i.e., of the object on a gray background at a canonical viewing angle), and the subject had to touch which of two images was in the preceding sample image. We used a two-alternative forced-choice stimulus-response paradigm—for a given task session, whether for marmoset or human, only a pair of objects was tested (e.g., one day would be camel-wrench, another would be wrench-leg). This was varied across sessions so that images of a given object were tested in all possible two-way task settings (e.g., camel vs. rhino, camel vs. wrench, and camel vs. leg). For a given behavioral session, each object was consistently on the same side of the decision screen (e.g., camel would always be on left and wrench always on right). One motivation for using this kind of two-way stimulus-response design is that it reduces the working memory demands on the participant. As soon as the participant recognizes what is in the image, they can plan their motor action. In part because of this simplicity, the two-way stimulus response paradigm is broadly used across the neurosciences, including in rodents^36^, and it therefore allows behavioral benchmarking for comparative studies across an array of candidate model species. Moreover, as reported in the main text, we found that the pattern of image-by-image errors was highly similar across humans who performed the two-way stimulus-response design and those that performed the match-to-sample design (the latter of which was collected in the previous work), in which pair-wise image discriminations were interleaved within a session rather than across sessions (r = 0.90; this fully-interleaved match-to-sample design was used in the previous work with the macaques). The similarity of these signatures across experimental designs suggests that when comparing model systems on purely perceptual grounds, a simplified, two-alternative stimulus-response task can serve as a universal method for comparative studies across animals—even those where a challenging match-to-sample task design may be difficult to employ.

For the rodent task, the design was similar to that of the core recognition task, but sample stimuli were not flashed and were instead present on the screen until the subject made their choice. We used this design because it mimicked Zoccolan and colleagues’ study^36^. Instead of requiring the marmosets to use a nose-poke into two ports, we had the marmosets touch one of two white circles on-screen, one on the left and one on the right, to indicate their choice. We used white circles rather than token images to avoid allowing the marmosets to employ a pixel-matching strategy with the image on the screen.

### Stimuli

For the core object recognition task in marmosets, we used a subset of images from a previous study in macaques (see Supplemental Figure 3 for all 400 images). We briefly describe the image generation process below, but see Rajalingham et al., 2018 for more details^43^. These stimuli were designed to examine basic-level^56^, core object recognition. They were synthetically generated, naturalistic images of four objects (camel, wrench, rhino, and leg) displayed on a randomly chosen natural image background. The four objects were a randomly selected subset of the twenty-four used in the previous work. The spatial (x,y) position of the center of the object, the two angles parameterizing its pose (i.e., 3d rotation), and the viewing distance (i.e., scale) were randomly selected for each image. Stimuli were designed to have a high degree of variation, with disparate viewing parameters and randomized natural image backgrounds, in an effort to capture key aspects of the challenge of invariant object recognition, and to remove potential low level confounds that may enable a simpler visual system, either biological or artificial, to achieve high performance^57^.

In addition to this main set of 400 test images (100 for each object), we employed four additional sets of images in training the marmosets to do the task (for training details, see “Marmoset training procedure” below). The first consisted of a single token image: the object rendered at the center of a gray background in a relatively canonical pose (e.g., side view of an upright camel). The second set consisted of 400 images of each object at random poses, positions, and scales, all on a uniform, gray background. The third set consisted of a different sample of images that were drawn from the same distribution of generative parameters as our 400 test images (i.e., variation in pose, position, and scale on randomly selected natural backgrounds). To assess the generality of our image-by-image performance scores, we collected marmoset behavior to the test images at two different sizes. Marmoset position and distance from the viewing screen was not strictly controlled but was nonetheless relatively stereotyped (see “Homecage testing boxes” below), and the two image sizes subtended ~11° and ~22° of visual angle. For humans, we collected a single image size. Humans were not constrained in how they held the tablets on which the images were displayed, and this image size subtended ~4-12° of visual angle, depending on how the tablet was held.

For comparing marmosets to rodents, identical images were used as in a prior study in rats; see Fig. 1a for all images, and see Zoccolan et al., 2009 for more details^36^. In brief: these stimuli were of one of two synthetic, artificial objects, which were rendered at the center of the screen on a uniform black background. Objects were rendered at one of nine different azimuthal rotations (−60° to +60°, in steps of 15°) and one of six different scales, spanning ~1.4 octaves of size (from 0.5x to 1.3x the size of a template; for marmosets, 1.0x subtended ~11 degrees visual angle). Rotations and scales were fully crossed for each object to yield a total of 108 images (9 rotations x 6 scales x 2 objects).

### Web-based, homecage behavioral training system

In part inspired by the high-throughput behavioral systems used in some rodent work^58,59^, we developed a system where behavior would be collected in parallel from a large number of animals.

#### Web-based behavioral platform

We developed the MkTurk web-based platform (*mkturk.com*) to collect the data on mobile devices (i.e., tablets) that could be deployed anywhere. We opted for a web-based system for a variety of reasons. First, MkTurk just needs a web browser to run, and as a result the setup and installation across tablets is relatively low cost, both in terms of the researcher’s time as well as money, as consumer touchscreen tablets are often cheaper than more traditional behavioral rigs. Second, such a system made it relatively turnkey to evaluate humans and marmosets in as similar environment as possible—we simply distributed the same tablets to humans, who performed the touchscreen tasks just as marmosets did. Third, being based on the web naturally enables real-time streaming of the animals’ performance to automatic analysis pipelines in the cloud, which allows seamless monitoring of the animals on any device that can access that web (e.g., a researcher’s smartphone). As a result, monitoring animal performance and troubleshooting issues in real-time is more straightforward. Moreover, since task parameters are passed from the cloud to the tablets in real time, task parameters can be adjusted on the fly remotely, which was particularly helpful during training of many subjects in parallel.

For external hardware, we leveraged the open-source Arduino platform (Arduino Leonardo) coupled to a low-power piezoelectric diaphragm pump for fluid delivery (Takasago Fluidic Systems) and RFID reader for individual animal identification (ID Innovations), all powered by a single 5V USB battery pack. Thus, our system may not be as powerful as some traditional, fully-equipped experimental rigs, but our behavior box was highly optimized for SWaP-C (size: ~1 ft^3^, weight: ~10 lbs, power: ~5W, and cost: < $1,000).

#### Marmoset homecage behavior collection

Marmosets were tested in an 8” x 9” x 11” (width x height x depth) modified nest box that was attached to their housing unit. They were granted access to this box three hours a day. Marmosets could move freely in and out of the testing box for *ad libitum* access to food during the three hours, and in some instances where RFID was employed for automated identification, marmosets could engage with cagemates in social behavior when not performing the task. Marmosets performed trials on touchscreen tablets (Google Pixel C) inserted into a vertical slot in the rear of the nest box. Once in the nest box, marmoset body and head position were not strictly controlled (e.g., via chairing or head-fixing), but marmosets were encouraged into a relatively stereotyped position in a few ways. First, the boxes were augmented with dividers on both the left and right sides to restrict degrees of lateral freedom while marmosets were performing the task. Second, access to the touch screen was only available through a small (3.5” by 1.5”) armhole in the front plexiglass barrier. Third, a metal reward tube was embedded in the front plexiglass barrier and positioned at the center of the screen. Given that marmosets tended to perform well, they received regular rewards of diluted sweetened condensed milk via this tube (1 part milk : 7 parts water), and they tended to position themselves at this tube consistently, yielding a relatively stereotyped viewing position (e.g., see Supplemental Movie 1). By collecting behavior from animals in their housing unit in the colony, we were able to acquire data from all marmosets simultaneously. Indeed, in part because of this high-throughput behavioral system, we were able to measure performance for each of 400 images with high precision by collecting many trials per image (trial split-half correlation for marmoset’s i1n, Pearson’s r = 0.94).

#### Human behavior collection

Human data was collected on the same tablet hardware using the same MkTurk web app software. As with the marmosets, humans were allowed to perform trials in their home environments. Moreover, we also measured human image-by-image performance with high precision and reliability (trial split-half correlation for human i1n, Pearson’s r = 0.91).

### Marmoset training procedure

Marmosets were trained on the task through a series of stages. The first stage aimed to familiarize the marmosets with the motor mapping. In this case the sample image that was flashed was simply one of the token images, and thus was the same as the images used as buttons in the decision stage. This task therefore required no visual processing beyond pixel-matching, but was helpful to familiarize marmosets with the task paradigm. In the second stage, we introduced random position, pose and scale, but still presented the images on plain gray backgrounds, and this task served as a helpful bridge between the simple motor-mapping component and the full-scale core object recognition task that was the goal of our training. In the third and final training stage, marmosets were trained on the high-variation images with disparities in position, pose, and scale and with natural-image backgrounds. When marmosets saturated performance on this final stage, we switched them to test images, and we report the behavior from these separate test images not used in the training stages.

### Behavioral metric: i1n

#### Definition of i1n

To compare human and marmoset core object recognition, we employed an image-by-image metric, the i1n, which has been shown in previous work to be highly discerning between visual systems^43^ and a useful tool for selecting behaviorally challenging stimuli for studying neural processing in ventral visual cortex^60^. The i1n measures the discriminability of each image, and is designed to be a metric of a system’s sensitivity, as formalized by signal detection theory. For each image *j*, the difficulty *V_j_* is defined as:

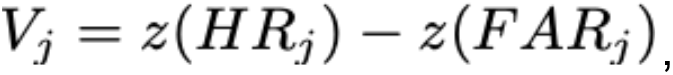

where *V* is a 400-length vector. *HR_j_* and *FAR_j_* are, respectively, the hit rate and the false alarm rate-the proportion of trials that image *j* was correctly classified and the proportion of trials that any image was incorrectly classified as that object. *z()* is the inverse cumulative density function of a Gaussian distribution, which allows evaluating the difference between the hit rate and false alarm in z-scored units. For images where a system was correct on every trial, the z-transformed hit rate is infinity, and we capped these values to be 4, as this value was just outside the ceiling of our empirically measured hit rates given the number of trials that were in each bin. No images for pooled marmosets nor humans reached this threshold; for individual human subjects (Fig. 3c), sub5 had 66 of 400 points at ceiling, sub0 had 8 of 400; excluding these ceiling points did not substantially alter the correlation between the two (all points: *r_nc_* = 0.861; excluding ceiling points: *r_nc_* = 0.871). Nine of the four hundred macaque images reached this threshold. We then subtracted off the mean value for each object to yield the i1n. Normalizing these values but subtracting off object-level performance (the “n” in the “i1n”), makes this metric robust to mean shifts in performance for each object and helps remove differential performance across behavior sessions. In the stimulus-responses design that we employed, object correlates with session. By contrast, when trials of different pairwise discriminations are interleaved within a session like in a match-to-sample design, any day-by-day variability in performance is not correlated with object-level performance. In practice, whether or not this object-level normalization was included did not affect the main results—Supplemental Figure 4 shows that key results are essentially the same when using an “i1” instead of an i1n.

#### Contextualizing the i1n

While the calculation of the metric was identical in our work and in the work by Rajalingham and colleagues^43^, differences in the inputs to the i1ns may lead to some mild differences between the nature of each of our i1ns. First, our i1n was computed with more images. Rajalingham and colleagues reported human-macaque correlations of image-level metrics on a subset of 240 of the 2400 images tested (10 of 100 images of each of the 24 objects), because human data was somewhat expensive to acquire and, image-level metrics require large amounts of data per image to be reliably estimated. In our work, because we concentrated data collection on four objects, we collected many trials on 100 images of each object, and thus computed i1n from 400 images total. The consequences of different numbers of images are likely not particularly substantial, though high correlation coefficients are somewhat less likely between 400-dimensional than with 240-dimensional vectors (e.g., the standard deviation of a null distribution of correlations coefficients between random Gaussian 240- and 400-dimensional vectors are, respectively, 0.065 and 0.050). A second, and potentially more consequential, difference between our i1n and the i1n used by Rajalingham and colleagues is that they measured the performance of each image against twenty-three different distractor objects, whereas we only measured ours against three. By averaging over far more distractor objects, their i1n likely minimizes the effect of the choice of distractor much more than our i1n does. Our i1n therefore likely measures something between Rajalingham and colleague’s i1n and their i2n, which averages over no distractors. Nonetheless, these differences between our i1n and the i1n used in previous work may not lead to substantial differences: as reported in the main text, we find that our 400-dimensional macaque i1n (averaged over three distractors) and their 240-dimensional macaque i1n (averaged over 23 distractors) are both equally similar to the corresponding human i1ns collected in comparable situations (*r_nc_*(macaque,human) = 0.77 in both cases).

#### Comparing i1ns: correcting correlation coefficients for test-retest reliability

To assess the similarity of different visual systems, we measured the consistency of the image-by-image performance of each by correlating i1ns (Pearson’s correlation coefficient). We subjected the “raw” correlation coefficients to a correction that accounts for the test-retest reliability of the data, as different systems’ i1ns will have different test-retest reliability due to, for instance, different amounts of data and1or different rates of errors. Reliability correction addresses potentially undesirable properties of comparing raw, uncorrected coefficients across pairs of systems. For instance, the consistency of a system with itself should be at ceiling (i.e., a correlation coefficient of 1), but an uncorrected correlation coefficient will have a value less than one, as it will be determined by the test-retest reliability. Because of this, the ceiling would in general be different across pairs of systems, and left uncorrected, this could lead to inaccurate inferences—e.g., one could naively conclude that system 1 and system 2 are more similar than system 1 and system 3, just because the researchers measured more reliable i1ns in system 2 than in system 3. To address these concerns, we measured and corrected for the test-retest reliability of the i1n for each system, by applying the correction for attenuation^61^, which estimates the noiseless correlation coefficient between the two—i.e., the correlation coefficient that would be observed as the number of trials goes to infinity. In doing so, we ensured that all comparisons between pairs of systems were on the same scale—i.e., the ceiling for each was indeed a correlation coefficient of 1—and thus were free to compare i1ns across these different systems.

We measured the noise-corrected correlation for a pair of systems’ i1ns by randomly partitioning trials for each image into two halves, computing i1ns for each half for each system, taking the mean of the correlation coefficients of i1ns across systems across the two split halves, and dividing it by the geometric mean of the reliability across systems (this denominator being the correction for attenuation):

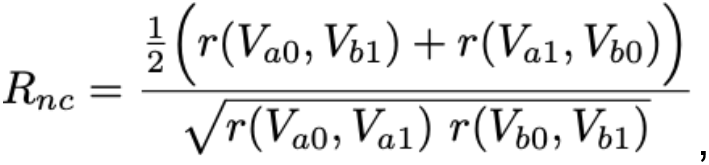

where *R_nc_* denotes the noise-corrected correlation coefficient, *r()* is a function that returns the Pearson correlation coefficient between its two arguments, *V_aO_* and *V_a1_* denote splits of trials for system a, and *V_bO_* and *V_b1_* denote splits of trials for system b. For each comparison, to mitigate variation due to how the data was randomly partitioned, we took the mean of this noise-corrected correlation coefficient across 1000 random partitions.

To test whether the correlation coefficients between two pairs of systems were different (e.g., marmoset-human correlation versus macaque-human correlation), we employed a dependent test of correlation coefficients^62,44^, which takes into account the dependency of the two correlation coefficients as they share a common variable (in this case humans). Accounting for this dependence increases the statistical power of the test.

Data analysis was conducted in Python and made use of the numpy^63^, scipy^64^, and sklearn^65^ libraries.

### Classifiers: Deep neural networks and pixels

#### Classifiers overview

To contextualize the behavior of simian primates, we also evaluated the overall performance and image-by-image performance for a variety of artificial systems. We trained linear classifiers for our task on top of representations from deep neural networks, which have been shown to be powerful models of visual behavior and neurophysiology^42,66,43^. We evaluated standard lmageNet-trained deep networks, downloading pretrained models from the torchvision package of PyTorch. As a low-level control, we also compared performance of linear classifiers trained on image pixels (256 x 256 pixel images), which in part assesses the extent to which features like luminance and contrast covary with labels and thus can be used to perform the task.

#### Classifiers for task performance

We evaluated linear classifiers on the penultimate layer of deep networks, training linear support vector machines (SVMs) on the same images and same binary tasks that the primates performed. We used a hinge loss and L2 regularization. The features were nonnegative, because they were the activations of rectified linear units in the network, and so we did not center the data as that would alter the consequences of L2 regularization We instead experimented with allowing the model to learn an intercept term or not, and observed similar results in both cases; we report results when learning an intercept. To select the strength of regularization, we searched over a range of 21 logarithmically spaced hyperparameters and selected hyperparameter values via 80-20 splits within the training set. To mitigate variation of results due to which images were selected for training or testing, we trained 1000 classifiers on random train-test partitions.

We confirmed that we were saturating performance with the amount of training data, by varying the number of images we trained on, using values of 10, 50, 90, and 99 images per object (out of 100 total images per object). Moreover, in a pilot experiment we established that we were not underestimating deep network performance due to the fact that the marmosets, humans, and macaques were able to generate an expanded “training set” of retinal activations because they were free to fixate anywhere during image presentation. To test this possibility, we generated an augmented training set that was 100x bigger, taking 100 random crops of different sizes and center locations (but not horizontal reflections) to mimic the varied fixations that marmosets and humans were allowed. We then trained classifiers on AlexNet representations, as that was the network with the lowest performance and thus greatest potential for improvement. lt seemed plausible that this augmented dataset would lead to improved performance, as convolutional networks are not invariant to modest translation or scaling^67^, and this kind of data augmentation is a central component of contemporary deep network training pipelines. Nonetheless, expanding the dataset hardly improved decoder performance at all, demonstrating that, at least for the amount of training data used here, varied “fixations” do not substantially improve classifier performance on top of a pretrained network, and thus we appear not to be overestimating the difficulty of this task for deep network representations.

#### Classifiers for i1ns

To evaluate the i1n of artificial visual systems (deep neural networks and pixel representations), we computed the distance of each image from the hyperplane learned by a linear SVM. We took 50-50 splits of images, trained classifiers on each half and evaluated distances for the left-out images. To derive an i1n for each artificial visual system, we averaged the distance for each image over its value in the three tasks, and subtracted off mean values for each task. To get highly reliable estimates of the network’s i1n, we performed this procedure 1,000 times for each task and each network, as well as for pixel representations. The ability to run arbitrarily large number of classifiers *in silico* yields the ability to drive the split-half correlation of the resulting i1ns arbitrarily high. Indeed, while the distances from the hyperplane are relatively reliable across individual partitions (for network classifiers, correlation coefficient ranges from 0.86 to 0.93; for pixels-based classifier: 0.66), the reliability of the distances averaged across 500 random train-test partitions is greater than 0.99 for all classifiers. Because of these exceedingly high test-retest reliabilities, we did not apply noise correction to the classifier i1ns learned from pixel or deep network features.

#### Classifiers for rodent task

To evaluate the kinds of visual representations required to generalize on the task used by Zoccolan and colleagues, we mimicked the training that the animals received in how we trained our classifiers. We trained classifiers on pixels from the 28 images used initially in training (14 images at either 0° rotation or 1.0x scale for each of two objects), and evaluated the performance of the resulting classifier on the 80 held-out test images (40 images for each of the two objects). We again used a linear SVM classifier with a hinge loss and L2 regularization and selected the regularization strength hyperparameter via cross-validation within the 28 train images. We evaluated 5 candidate values for regularization strength which were logarithmically spaced, and performed 10 random splits within the training set, training classifiers with each regularization strength on 23 images and evaluating the quality of the fit with the remaining 5. We then selected the regularization strength that performed best on left-out train images, and trained a single classifier with all 28 images with this regularization coefficient. We evaluated the performance of this classifier on the unseen 80 test images, and found that it classified all but two correctly (overall performance: 97.5%)-the two that it got incorrect were at the smallest size (0.5x) combined with the most dramatic rotation (60° or −60°) (bottom left and right corners of Fig. 3a). Given the already high performance of image pixels, we did not further evaluate deep network performance on this task (but see Vinken and Op de Beeck^68^).

## Acknowledgements

The authors thank Robert Desimone, James DiCarlo, and Guoping Feng for early project support; Hector Cho, Elizabeth Yoo, and Michael Li for technical support; James DiCarlo and Rishi Rajalingham for the macaque data; Davide Zoccolan for providing the original images used in the rat behavior study; and Hector Cho, Aniruddha Das, Asif Ghazanfar, Nancy Kanwisher, Jack Lindsay, Rishi Rajalingham, and Erica Shook for comments on a previous version of the manuscript. The work was funded by an NlH Postdoctoral NRSA fellowship to A.K. (F32 DC017628), an NlH R00 to E.l. (EY022671), and a Klingenstein-Simons Fellowship in Neuroscience to E.l.

## Author Contributions

A.J.E.K. and E.B.l designed the research. A.J.E.K., S.L.B., Y.J, T.T., and E.B.l. developed the experimental apparatus. A.J.E.K., S.L.B., Y.J., T.T., and E.B.l. collected the data. A.J.E.K. analyzed the data. A.J.E.K. and E.B.l. wrote the manuscript. All authors reviewed and edited the manuscript. E.B.l. acquired funding and supervised the research.

---

**Supplemental Figure 1.**
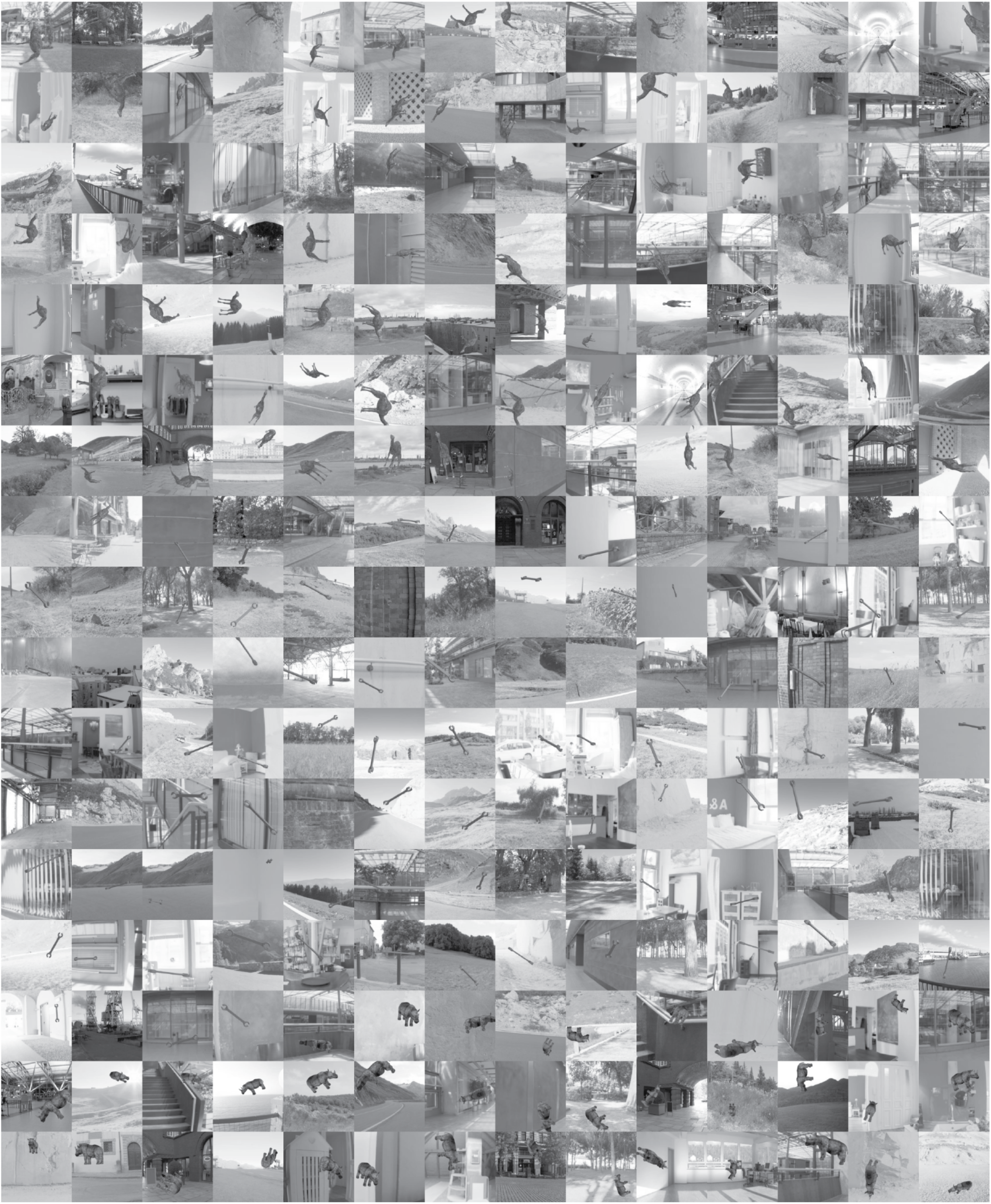
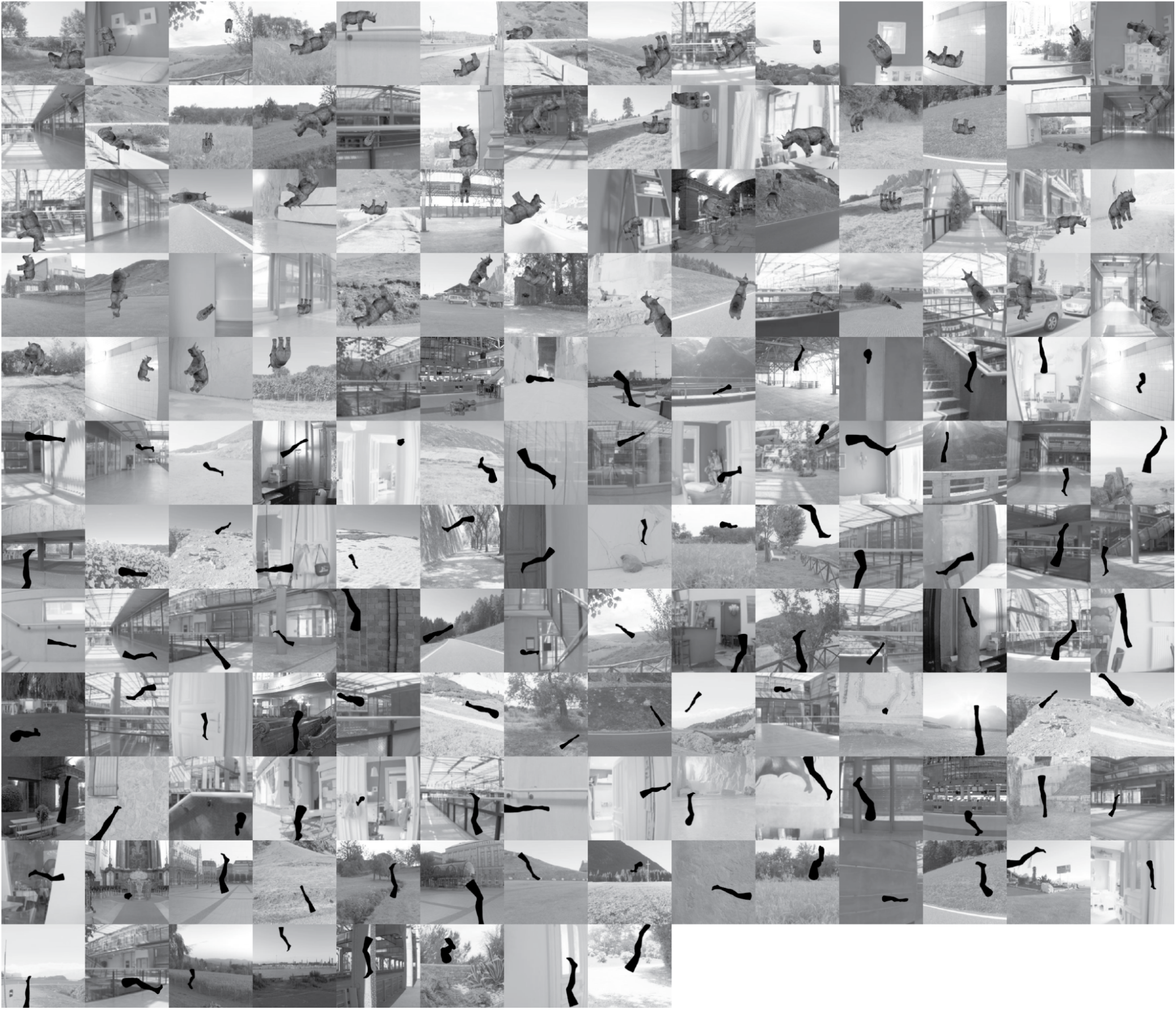
All 400 images. All four hundred images of camels, wrenches, rhinos, and legs superimposed on natural backgrounds. We measured marmoset and human performance on each of these images, and compared these performances with each other and with macaque performance on the same images collected in previous work^22^.

**Supplemental Figure 2.**
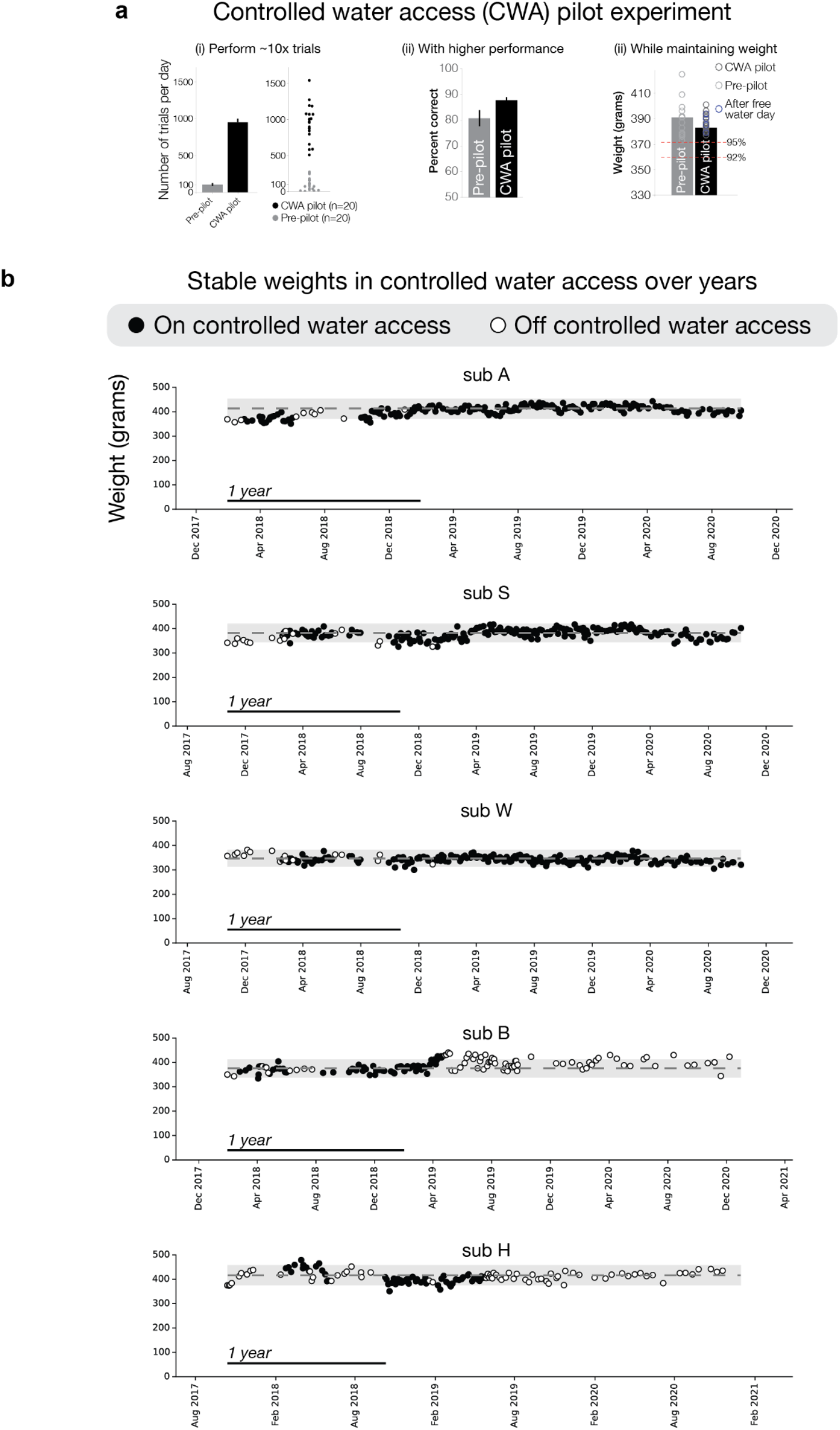
Controlled water access is safe and effective in marmosets over extended periods of time. a. **Controlled water access pilot experiment**. In controlled water access paradigms, animals only receive fluid from performing the task. This approach, though standard in macaque and rodent work, had been largely avoided in marmosets, potentially due to a reputation of being too fragile. We evaluated the potential of using water restriction in marmosets, proceeding carefully at first with a small pilot experiment designed to test whether controlled water access may be safe and effective. We directly compared behavior of one animal when measured under controlled water access versus *ad libitum* water access. In an effort to give the *ad libitum* condition the greatest chance of success, in this condition we removed the marmoset’s access to water three hours before the task to induce a greater probability of thirst and used a high-value reward during the task (sweetened condensed milk). Nonetheless, under the controlled water access condition, the marmoset performed approximately 10x the number of trials (i), with higher performance (ii), while maintaining a relatively stable weight (iii). b. **Stable weights in chronic controlled water access for more than a year**. In follow-up studies using more animals and longer time periods, animals maintained stable weights under controlled water access for more than a year. Dashed line indicates baseline weight measured before CWA. Shaded area is that baseline +/−10%.

**Supplemental Figure 3.**
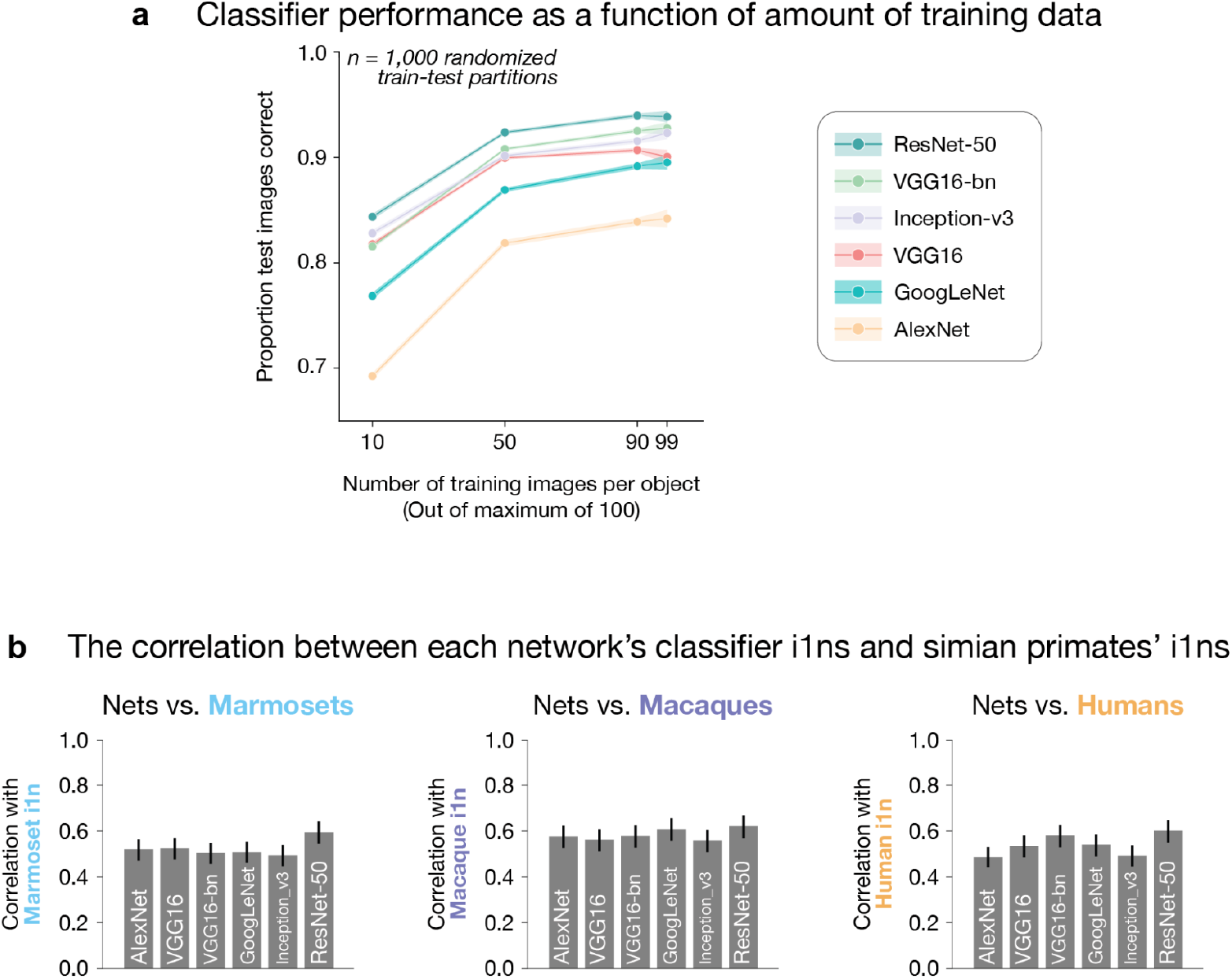
Deep networks: Performance as a function of amount of training data and similarity to simian primates in i1n. a. **Deep network performance on core recognition task**. The difficulty of a discrimination task varies drastically depending on the choice of stimuli^21^, and so we verified that our images yielded a challenging task not only by assessing human performance as we report in the main text, but also by evaluating the performance of state-of-the-art engineering systems. We trained binary classifiers atop the penultimate layer of artificial deep neural networks, using 10, 50, 90, or 99 images per object (total images per object: 100), varying the training regime to establish that we reached a performance plateau with the amount of training data that we had available. Each line is a different network (VGG16-bn denotes a VGG16 architecture trained with batch normalization). Error bars are SEM over 1,000 classifiers trained on random train-test partitions; standard errors increase as amount of training data increases since the test dataset size concomitantly decreases (i.e., train size of 99 leaves only 1 image per object for testing). As we report in the main text, the raw input was insufficient to support task performance, as image pixel representations performed near the 50% chance level. Additional sensory processing, as instantiated by deep artificial neural networks, yielded performance at 84-94% indicating that high performance was achievable, but that even high-quality computer vision systems did not readily perform the task perfectly. These analyses complement the human performance results in demonstrating that recognizing the objects in these images requires nontrivial sensory computation. b. **Correlation between classifiers on different networks’ features and simian primates**. From left to right, we compared marmoset, macaque, and human i1ns with i1n of the six networks. The consistency between networks and simians was relatively similar across the different networks.

**Supplemental Figure 4.**
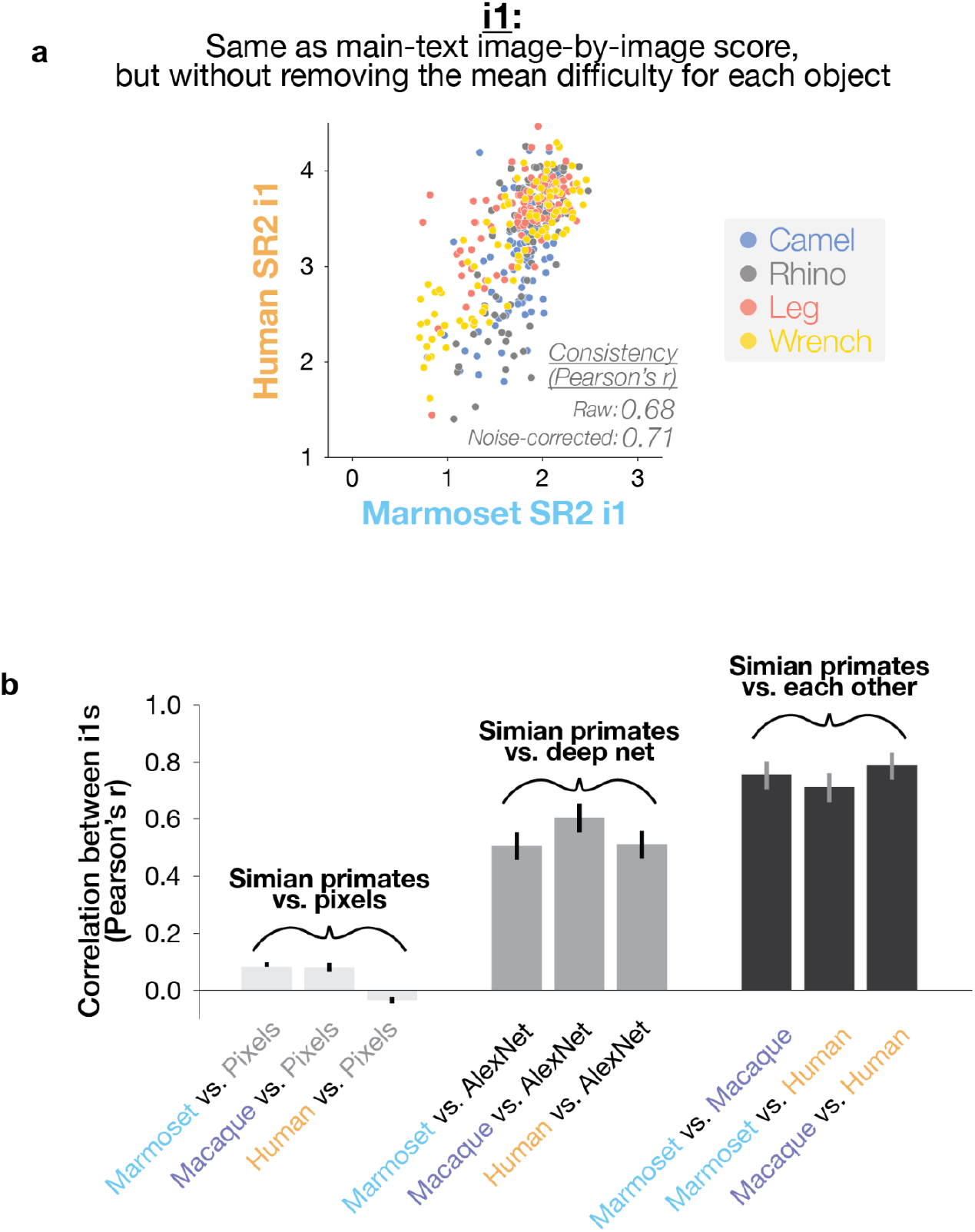
Image-by-image behavioral similarity between marmosets and humans is robust to details of the comparison metric. **a.** The image-by-image difficulty score in the main text (i1n) had the mean difficulty of the object removed from each image-wise score (see main text Fig. 2a). Previous work had found that such a metric of within-object image difficulty is highly discriminating between model systems (Rajalingham et al., 2018). Here, we plot the same image-by-image metric, but without this object-level de-meaning step. We find that that the similarity between marmosets and humans is robust to this detail of the comparison metric. Additionally, as this metric does not include a de-meaning step, this plot shows the difference in overall performance level between marmosets and humans-marmoset image-wise d’ ranges from just under 1 to approximately 2.5, while human d’ scores range from just under 2 to just over 4. While overall performance varies across simians (as shown in main text Fig. 1e), the comparative difficulty of each image was relatively robust as seen in the high correlation between marmosets and humans (here and main text, Fig. 2). b. When we compare the unnormalized i1s from pixel classifiers, deep network classifiers, and simian primates, we see a similar pattern of results as we see with i1ns (compare to main text, Fig. 2g).

